# A GATA factor radiation in *Caenorhabditis* rewired the endoderm specification network

**DOI:** 10.1101/2022.05.20.492851

**Authors:** Antonia C. Darragh, Rachel Weinstein, Jessica R. Bloom, Scott A. Rifkin

## Abstract

Although similar developmental regulatory networks can produce diverse phenotypes, different networks can also produce the same phenotype. In theory, as long as development can produce an acceptable end phenotype, the details of the process could be shielded from selection, leading to the possibility of developmental system drift, where the developmental mechanisms underlying a stable phenotype continue to evolve. Many examples exist of divergent developmental genetics underlying conserved traits. However, studies that elucidate how these differences arose and how other features of development accommodated them are rarer. In *Caenorhabditis elegans*, six transcription factors that bind motifs with a GATA core sequence (GATA factors) comprise the zygotic part of the endoderm specification network. Here we show that the core of this network - five of the genes - originated within the genus during a brief but explosive radiation of this gene family and that at least three of them evolved from a single ancestral gene with at least two different spatio-temporal expression patterns. Based on analyses of their evolutionary history, gene structure, expression, and sequence, we explain how these GATA factors were integrated into this network. Our results show how gene duplication fueled the developmental system drift of the endoderm network in a phylogenetically brief period in developmentally canalized nematodes.

## Introduction

Six of the 11 GATA factors in *C. elegans* function in the endoderm specification network (Fig. 1; McGhee 2013; Maduro 2017). In this network, the maternal transcription factor SKN-1 initiates a feedforward cascade in which GATA factors that specify, differentiate, and maintain the cell fate of the endoderm are expressed (Bowerman et al. 1992; Blackwell et al. 1994; Maduro and Rothman 2002). SKN-1 is a basic leucine zipper (bZIP)/homeodomain-like transcription factor (Bowerman et al. 1992) that first activates transcription of the functionally redundant GATA factors *med-1* and *med-2* in the endomesoderm (EMS) cell (Fig. 1; Maduro et al. 2001). MED-1, MED-2, and possibly SPTF-3 (Sullivan-Brown et al. 2016), a specificity protein transcription factor, activate expression of the largely functionally redundant GATA factors *end-3* and then *end-1* during the subsequent two cell divisions in the endoderm lineage (E and 2E stages) (Fig. 1; Maduro and Rothman 2002; Baugh et al. 2003; Maduro, Hill, et al. 2005; Maduro et al. 2015). POP-1 and PAL-1 are other maternally provided transcription factors (HMG box and homeoprotein, respectfully) (Lin et al. 1995; Hunter and Kenyon 1996) that make a minor contribution to endoderm specification (Fig. 1). In fact, only when other parts of the network are disrupted does knocking out *pal-1* show an effect (Maduro et al. 2001; Maduro, Hill, et al. 2005; Maduro et al. 2007; Maduro et al. 2015). Wnt/MAPK-induced POP-1 and SYS-1, a beta-catenin cofactor, together likely directly activate *end-1* expression (Shetty et al. 2005; Phillips et al. 2007). *C. elegans* SKN-1 binding sites (Blackwell et al. 1994) are also found in within X bp upstream of most *Caenorhabditis end-3* and *end-1* predicted coding sequences suggesting that SKN-1 also directly activates them (Zhu et al. 1997; Maduro, Kasmir, et al. 2005a; Maduro 2020). END-3 and END-1 then activate the expression of the GATA factors *elt-7* and *elt-2* in the 2E (when there are two endodermal cells)and 4E (when there are four endodermal cells) stages, respectively (Zhu et al. 1998; Maduro and Rothman 2002; Sommermann et al. 2010; Boeck et al. 2011) (Fig. 1). ELT-7 and ELT-2 are partially redundant in regulating and directing the differentiation and maintenance of the endoderm from the 2E stage to the final twenty intestinal cells that comprise the entire endoderm of these worms (Sulston et al. 1983; Fukushige et al. 1998; Fukushige et al. 1999; McGhee et al. 2007; McGhee et al. 2009; Sommermann et al. 2010; Dineen et al. 2018). *elt-4*, a likely degenerate duplicate of *elt-2,* is expressed later in the development of the endoderm but does not have any known function (Fukushige et al. 2003). The four other canonical GATA factors in *C. elegans* (*elt-3, elt-1, elt-6,* and *egl-18*) all function during hypodermal (ectoderm) cell development (Gilleard and McGhee 2001; Koh and Rothman 2001).

**Figure 1.**
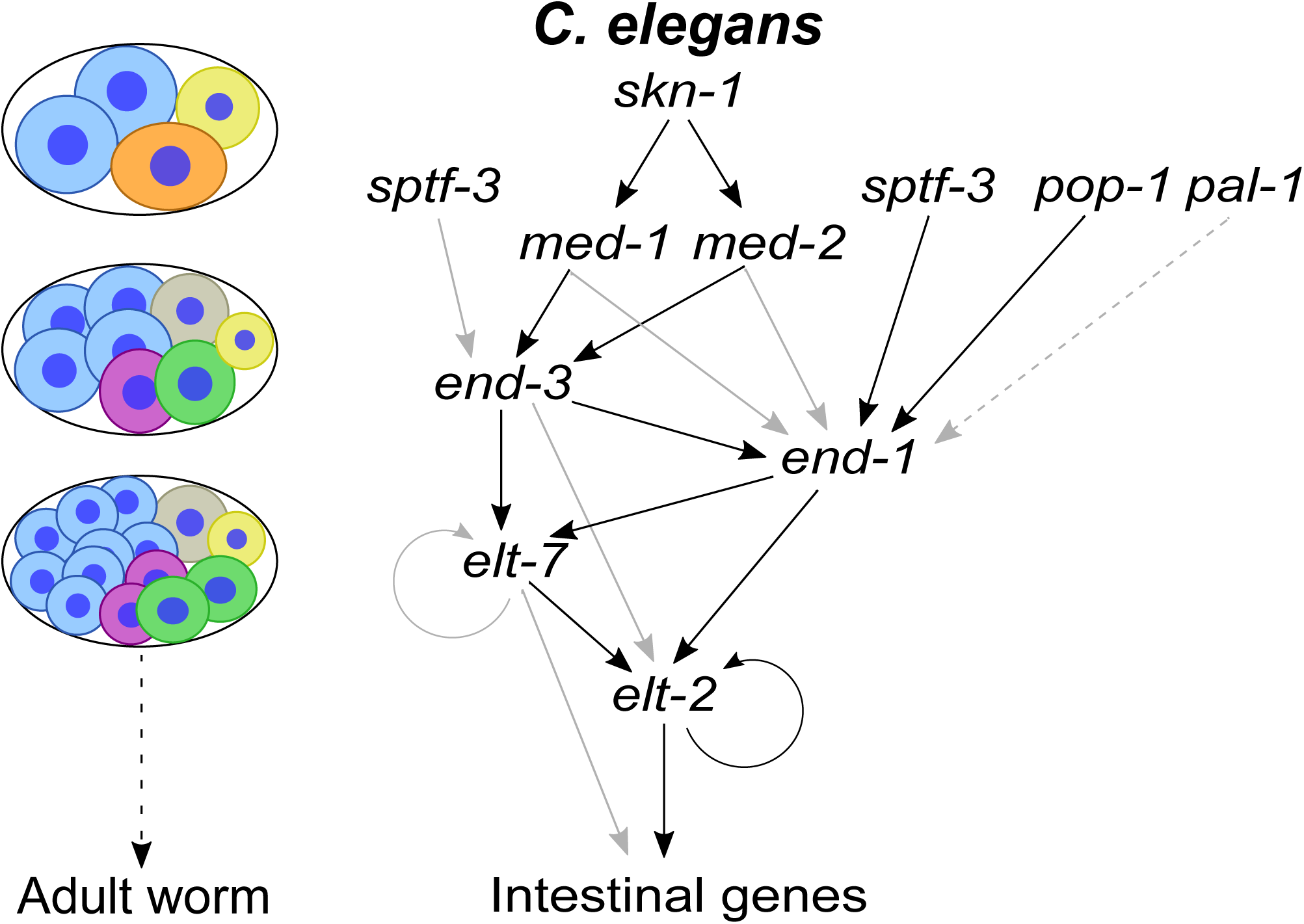
*C. elegans* endoderm specification network. The *C. elegans* endoderm specification network is shown on the right and the approximate embryonic stages during which most of the gene expression associated this network takes place is shown on the left. This network is initiated primarily by SKN-1 in the EMS cell (orange cell on bottom of four-cell embryo); however, SPTF-3, POP-1, and possibly PAL-1 also contribute to the activation of this GATA factor cascade, as shown. Six of the 11 *C. elegans* GATA factors (*med-1*, *med-2*, *end-3*, *end-1*, *elt-7*, and *elt-2*) function in this network (as shown). *med-1* and *med-2* expression initiates in the EMS cell and MED-1 and MED-2 regulate genes in both the first endoderm (1E) cell (green cell on bottom right of eight-cell embryo) and the first mesoderm (MS) cell (purple cell to the left of the 1E cell). *end-3* expression starts in the late EMS or early 1E cell while *end-1* expression starts in the late 1E or early two endoderm (2E) cell stage (two green cells in 14-cell embryo). *elt-7* expression starts in the 2E cells and *elt-*2 expression starts near the beginning of the 4E cell stage (not shown). Black arrows indicate well supported regulatory connections, while gray and dashed gray arrows represent weaker and not as well supported interactions, respectively.

GATA factors are potent endodermal regulators throughout bilaterians (Patient and McGhee 2002; Gillis et al. 2007) even when expressed heterologously. For example, if *C. elegans* END-1 is expressed in explant *Xenopus laevis* animal caps (an ectodermal lineage) it activates endoderm development, demonstrating both conservation of endoderm specification capabilities between ecdysozoa and vertebrates and a surprising generalized ability of this GATA factor to function despite a markedly different intracellular context (Shoichet et al. 2000). In *C. elegans*, using the *end-1* promoter to highly express ELT-2 or ELT-7 can compensate for the loss of all four of the genes *end-3*, *end-1*, *elt-7*, and *elt-4* (Wiesenfahrt et al. 2016)suggesting that the primary functional difference among these endoderm-specific GATA factors is not in their ability to bind DNA and regulate downstream genes but rather in when they are expressed.. However, expression of neither *C. elegans elt-3*, which encodes a hypoderm-specifically expressed GATA factor, nor *Mus musculus* GATA-4 expressed using the same *end-1* promoter, were able to rescue loss of both *end-1* and *end-3* in *C. elegans* (Wiesenfahrt et al. 2016), suggesting that GATA factors are not all interchangeable. Identifying attributes responsible for functional redundancy among some GATA factors has been difficult because these proteins have diverged extensively outside of their DNA-binding domains (Lowry and Atchley 2000; Gillis et al. 2008; Eurmsirilerd and Maduro 2020; Maduro 2020; see below) and (other than the MED orthologs (Broitman-Maduro et al. 2005; Lowry et al. 2009; Eurmsirilerd and Maduro 2020; see below) they are all thought to bind to canonical HGATAR DNA sites (Gerstein et al. 2010; Araya et al. 2014; Narasimhan et al. 2015; Du et al. 2016; Wiesenfahrt et al. 2016; Maduro 2020; see below).

Over the last 50 years, many studies have demonstrated that gene duplication is a major mechanism through which new genes with novel functions evolve (e.g., Ohno 1970; Gottlieb 1977; Escriva et al. 2006; Assis and Bachtrog 2013; McKeown et al. 2014). Four possible models of paralog divergence currently dominate the literature: pseudogenization (Ohno 1970; Nei and Roychoudhury 1973; Charlesworth et al. 1994; Lynch and Walsh 1998; Eyre-Walker and Keightley 1999; Denver et al. 2004; Haag-Liautard et al. 2007), neofunctionalization (Ohno 1970), subfunctionalization (Hughes 1994; Force et al. 1999; Lynch and Force 2000), and redundancy (Ohno 1970; Nei et al. 2000; Piontkivska et al. 2002; Kondrashov and Kondrashov 2006). Evidence for each of these evolutionary outcomes of gene duplication can be found in nature (e.g., Jozefowicz et al. 2003; He and Zhang 2005; Gout and Lynch 2015), but it is often difficult to determine the exact evolutionary trajectory since information on extant paralogs is often compatible with several possible histories and these categories are not necessarily mutually exclusive for specific paralog pairs (Gera et al. 2022).

A recent comparison of nematode GATA factors found that the *elt-3* family of orthologs had undergone the most gene duplications and sequence divergence, suggesting that this gene may have evolved faster than the other GATA factors in the phylum (Eurmsirilerd and Maduro 2020). Maduro (Maduro 2020) found that orthologs of five of the six GATA factors that regulate endoderm development in *C. elegans* – *med-1*, *med-2*, *end-1*, *end-3*, and *elt-7* – are specific to the *Elegans* supergroup, suggesting that these genes arose in the ancestor of this clade. Maduro proposed a model for the origin of the regulatory network specifying endoderm in *C. elegans* based on analysis of a subset of *Caenorhabditis* GATA factors in the genomes of 20 species in the *Elegans* supergroup and four species from outside of that supergroup. We took advantage of draft sequences of the genomes of an additional 34 *Caenorhabditis* species (Stevens 2020) to carry out a more comprehensive analysis of GATA factors throughout the *Caenorhabditis* genus and determine the origin of the *C. elegans*-type endoderm specification network.

To identify GATA factors in *Caenorhabditis*, we searched for their characteristic DNA-binding domain in all fifty-eight *Caenorhabditis* species for which genomic sequence assemblies or transcriptomes were available and in the genomes of two outgroup *Diploscapter* species. We estimated the evolutionary history of this gene family using maximum likelihood approaches. We focus here on the effects of an *elt-3* radiation on the developmental genetics of endoderm specification. This study illustrates how gene duplications fueled the evolution and elaboration of an essential regulatory network, all without causing any obvious change in development or morphology.

## Results

The preponderance of genes from species in the *Elegans* and *Japonica* groups in our GATA-domain containing phylogeny (Fig. 2) indicates that the GATA factor family dramatically radiated within the *Elegans* supergroup, expanding from a typical six genes per species to at least 10 (median of 16, maximum of 34). This radiation occurred along two consecutive internal branches in the species tree (Fig. 2), suggesting a concerted burst of gene duplication that affected different developmental genetic networks, including the endoderm specification network^26^, during a phylogenetically brief period. In this paper we focus on the expansion highlighted by the black arrow in Figure 2B; analyses of the other expansions will be published elsewhere.

**Figure 2.**
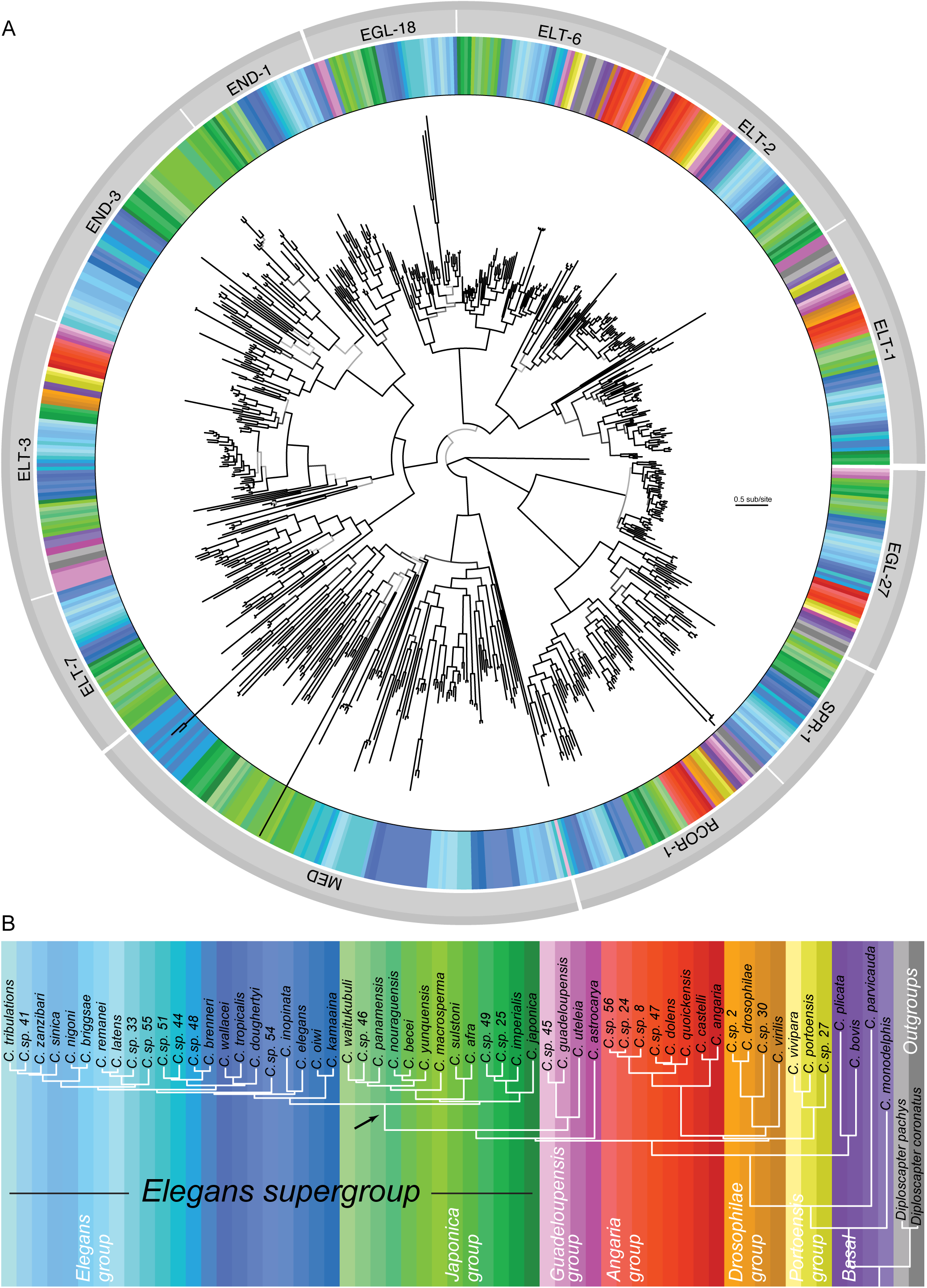
Inferred evolutionary history of *Caenorhabditis* GATA-domain-containing proteins. **(A)** Maximum likelihood phylogeny of 714 “confident” GATA-domain-containing proteins in 58 *Caenorhabditis* and two outgroup nematode species. A GATA factor from the slime mold *Dictyostelium fasciculatum* was used to root the phylogenic tree (located between the ELT-1 and EGL-27 ortholog groups). The tree includes both canonical GATA factors and EGL-27, SPR-1, and RCOR-1 orthologs which are proteins that contain atypical GATA-binding domains but which scored above our threshold on the PROSITE GATA-type ZnF domain profile. The colors in the ring encircling the tree correspond to the species in which the protein was identified (the key to color-species correspondence is given in C below). The names of the 12 ortholog groups the 714 proteins were categorized into are indicated in the lighter of the two outer gray rings (with white gaps between groups). Clades comprising multiple ortholog groups are highlighted by the darker gray outer ring (with white gaps between clades). The intensity of shading of each branch of the tree is indicative of its degree of bootstrap support, darker shading indicates stronger support. The key for translating branch length into evolutionary distance (in units of amino acid substitutions per site) is shown to the right of the tree. **(B)** Phylogenetic relationships among the 60 species used in this study (based on Stevens (2020)). Each species is designated by a different color shade; color-species designations are the same as used in (B) above. The black arrow points to the *Elegans* supergroup ancestral branch where the ancestral *med*, *end-1*, *end-3*, and *elt-7* genes, as we know them from *C. elegans*, likely arose.

Of the endoderm network GATA factors, ELT-2 is found in every species in the genus, but MED, END-3, END-1, and ELT-7 orthologs are absent from non-*Elegans* supergroup species (Fig. 2; see also Eurmsirilerd and Maduro 2020). This suggests a perplexing developmental question: how do these species specify endoderm when they are missing the genes that comprise the central part of the endoderm specification network that we know about from *C. elegans*? To answer this question, we started by investigating whether the role of ELT-2 in endoderm development is conserved.

### The role of ELT-2 is likely conserved throughout the genus

Using single molecule fluorescence *in situ* hybridization (smFISH) (Raj et al. 2008) we examined *elt-2* expression in *C. angaria*, of the *Angaria* group. *C. angaria elt-2* expression resembles that of *C. elegans* (Fig. 3A,C,D); i.e., its expression is restricted to the endoderm, starts in the 4E cell stage, and continues throughout embryonic development. Data from RNA-sequencing in *C. angaria* (Macchietto et al. 2017) corroborates this expression pattern).

**Figure 3.**
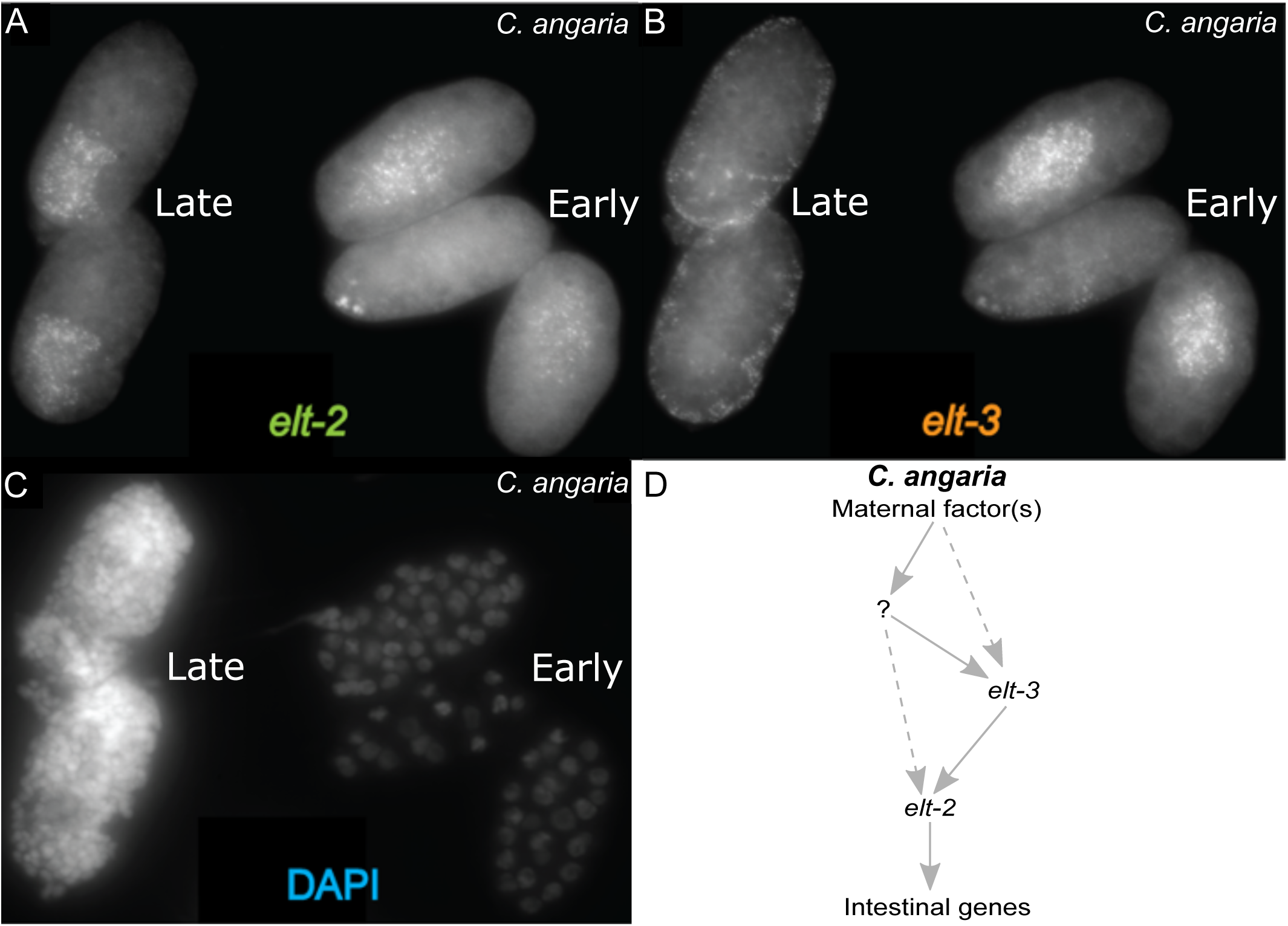
Expression of *elt-3* and *elt-2* mRNA in *C. angaria*, a non-*Elegans* supergroup species. **(A-C)** Image of five embryos, each at a different developmental stage, illustrating the patterns of *elt-3* and *elt-2* mRNA expression observed in *C. angaria* using smFISH. The embryo depicted at the top left is at the comma stage (approximately) and contains more than 100 cells; the embryo at the bottom left is at the bean stage (approximately) and contains more than 100 cells; the embryo at the top right contains 54 cells; the embryo in the middle on the right contains 16 cells; and the embryo at the bottom right contains 25 cells. **(A)** Visualization of *C. angaria elt-2* mRNA after hybridization with a smFISH probe specific for *C. angaria elt-2*. **(B)** Visualization of *C. angaria elt-3* mRNA after hybridization with a smFISH probe specific for *C. angaria elt-3*. **(C)** DAPI-stained nuclei of *C. angaria* embryos (proxy for developmental stage). **(D)** Model of *C. angaria* endoderm specification network based on these smFISH results.

*C. elegans* ELT-2 prefers to bind TGATAA sites. For example, McGhee and colleagues (McGhee et al. 2007; McGhee et al. 2009) identified 197 genes in *C. elegans* expressed specifically or predominantly in the intestine and found that the putative promoters of these genes are enriched with TGATAA sites (McGhee et al. 2007; McGhee et al. 2009). Analysis of one of these genes showed that *C. elegans* ELT-2 interacts with these sites *in vivo* to regulate its targets(Lancaster and McGhee 2020). Using reciprocal BLASTp (Altschul et al. 1990; Camacho et al. 2009), we identified orthologs of these *C. elegans* intestine-expressed genes in 57 other sequenced *Caenorhabditis* species and found that many of their putative intestinal promoters are also enriched for TGATAA sites compared to promoters for orthologs of muscle, hypoderm, and neural genes (Fig. 4).

**Figure 4.**
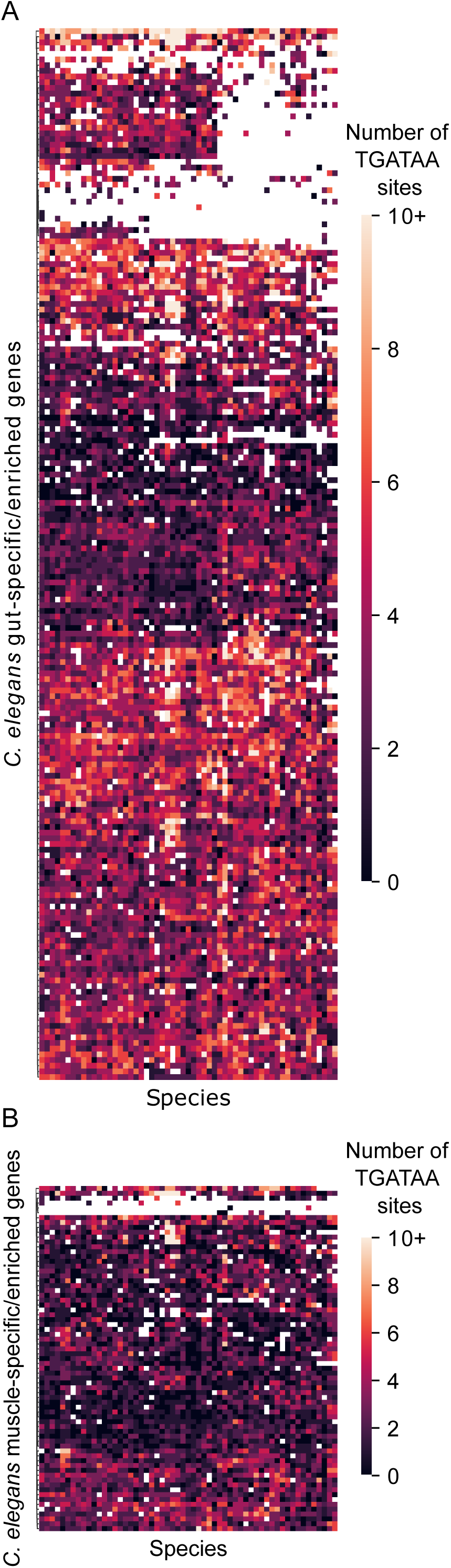
Conservation of TGATAA sites in putative promoters of orthologs specifically expressed or enriched for expression in gut and muscle. Heatmaps of the number of TGATAA sites in the promoter regions of orthologs expressed specifically or primarily in (A) gut versus (B) muscle in the 59 non-*C. elegans* species included in this study. The columns comprising the x-axis represent each species, in the same order (left to right) as the listing of species in the phylogeny shown in Figure 2B. Each row on the y-axis represents the promoter region of a *C. elegans* gene (McGhee et al. 2007; McGhee et al. 2009), ordered using hierarchical clustering with Euclidean distance metric. The color key is shown to the right of each heatmap plot. To make the color scaling more informative, the few promoter regions that had more than 10 TGATAA sequences are shown as having only 10 TGATAA sites within their promoters. White space in heatmaps indicates species for which we did not find an ortholog for that *C. elegans* gene. **(A)** Promoters of *C. elegans* orthologs specifically expressed or enriched for expression in gut. **(B)** Promoters of *C. elegans* orthologs specifically expressed or enriched for expression in muscle.

### ELT-3 is the closest relative to END-1, END-3, and ELT-7 and has a broader expression pattern outside of the Elegans supergroup

Even if ELT-2 organizes endoderm development in non-*Elegans*-supergroup species, its expression starts too late for it to be activated by maternal factors. There must be an intervening gene (or genes) that transmits and refines the positional signal from maternal factors (Wagner 2014) over several cell divisions. This is the role played by the MED-1, MED-2, END-3, END-1, and ELT-7 GATA factors in *C. elegans* (Fig. 1). Our GATA factor phylogeny presents a clue to the possible identity of one such intervening gene: since END-3, END-1, and ELT-7 orthologs group together with ELT-3 orthologs in a well-supported clade (Fig. 2A), *elt-3* might fill that role.

In *C. elegans*, ELT-3 is expressed only in differentiating and differentiated hypoderm cells (Gilleard et al. 1999; Gilleard and McGhee 2001), which makes it an unlikely candidate for involvement in endoderm development. Indeed, even when expressed under the control of the *C. elegans end-1* promoter, in the right place and at the right time (in a *C. elegans end-1* and *end-3* double mutant), *C. elegans elt-3* is unable to initiate endoderm specification (Wiesenfahrt et al. 2016), despite being able to bind to TGATAA sites *in vitro* (Narasimhan et al. 2015). However, unlike its paralogs, ELT-3 is found in all species in the genus raising the possibility that its role in *C. elegans* is not indicative of its ancestral function.

To investigate ELT-3’s role in non-*Elegans*-supergroup species we visualized *elt-3* expression in *C. angaria* embryos using smFISH (Raj et al. 2008). We found that *C. angaria elt-3* is expressed in two phases. The first phase starts at the 2E stage with expression restricted to the endoderm which erodes after the 4E cell stage (Fig. 3B-D), timing that resembles that of *C. elegans end-1* (Raj et al. 2010). Moreover, single-embryo RNA-sequencing revealed expression of *elt-3* in *C. angaria* that was slightly earlier and at higher levels than in *C. elegans* (Macchietto et al. 2017) and a blip of higher expression in *C. angaria* that coincides with the timing of *end-1* expression in *C. elegans*. In the second phase, *elt-3* in *C. angaria* is not expressed in endodermal but in hypodermal cells (Fig. 3B,C), resembling *elt-3* expression in *C. elegans* (Gilleard et al. 1999; Gilleard and McGhee 2001).

### The organization of the elt-2 promoter differs markedly between Caenorhabditis subclades although regulation by a HGATAR-binding transcription factor is likely conserved throughout the genus

To investigate how *elt-2* was regulated before the *elt-3* subclade expansion (see above), we searched for conserved transcription factor binding sites in the *elt-2* promoters of non-*Elegans* supergroup species (see Materials and Methods). We found significantly (p-value less than 0.05) more canonical GATA factor binding sites, i.e. HGATAR (Ravagnani et al. 1997); in all but two of these *elt-2* promoters than expected by chance (Fig. 5A). We also examined *Elegans* supergroup *elt-2* promoters and found a striking conservation of HGATAR sites in them (Fig. 5A). There are six highly conserved HGATAR sites in most *Elegans* supergroup *elt-2* promoters (highlighted by the colored dots at the top of Fig. 5A). Three of these sites comprise the sequence TGATAA in all *Elegans* supergroup species, the only exception being *C. elegans* which does not have an HGATAR site that aligns with the most 3’ of these three sites (sites under the pink dots in Fig. 5A). TGATAA sites are important for *C. elegans elt-2* expression (McGhee et al. 2007; McGhee et al. 2009; Du et al. 2016; Wiesenfahrt et al. 2016). They comprise the most overrepresented transcription factor binding site in *C. elegans elt-2* target genes(McGhee et al. 2007; McGhee et al. 2009), and TGATAA sites have been found to be the sites that *C. elegans* ELT-7, ELT-6, and ELT-3 GATA factors prefer to bind to (Narasimhan et al. 2015). Moreover, *C. elegans* ELT-2, ELT-7, END-3, and END-1 bind to TGATAA sites *in vitro* (McGhee et al. 2007; McGhee et al. 2009; Du et al. 2016; Wiesenfahrt et al. 2016). Two other conserved HGATAR sites, AGATAG and CGATAA, are found in all *Elegans* supergroup *elt-2* promoters (sites under the yellow and blue dots, respectively, in Fig. 5A). The sixth HGATAR site is the least well conserved; in 23 of 35 *Elegans* supergroup species CGATAG comprises this site, but in four species its sequence is AGATAA, in three species TGATAG, in two species AGATAG, and three species do not have a conserved HGATAR site that aligns at this position (see under the green dots in Fig. 5A). A few of these HGATAR sites similarly align in some non-*Elegans* supergroup species. However, no non-*Elegans* supergroup species contains more than one of these six sites (Fig. 5A). Overall, HGATAR sites are less abundant and less spatially conserved in the promoters of *elt-2* orthologs in non-*Elegans* supergroup species as compared to *elt-2* promoters in *Elegans* supergroup species (Fig. 5A). The organization of the *elt-2* promoter in *Elegans* supergroup species evolved in parallel with the expansion of GATA factors involved in the endoderm specification network (see above) and has remained highly conserved since. We found no evidence of overrepresentation of non-GATA-factor-binding sites among the non-*Elegans* supergroup *elt-2* promoters we analyzed (Fig. 5A).

**Figure 5.**
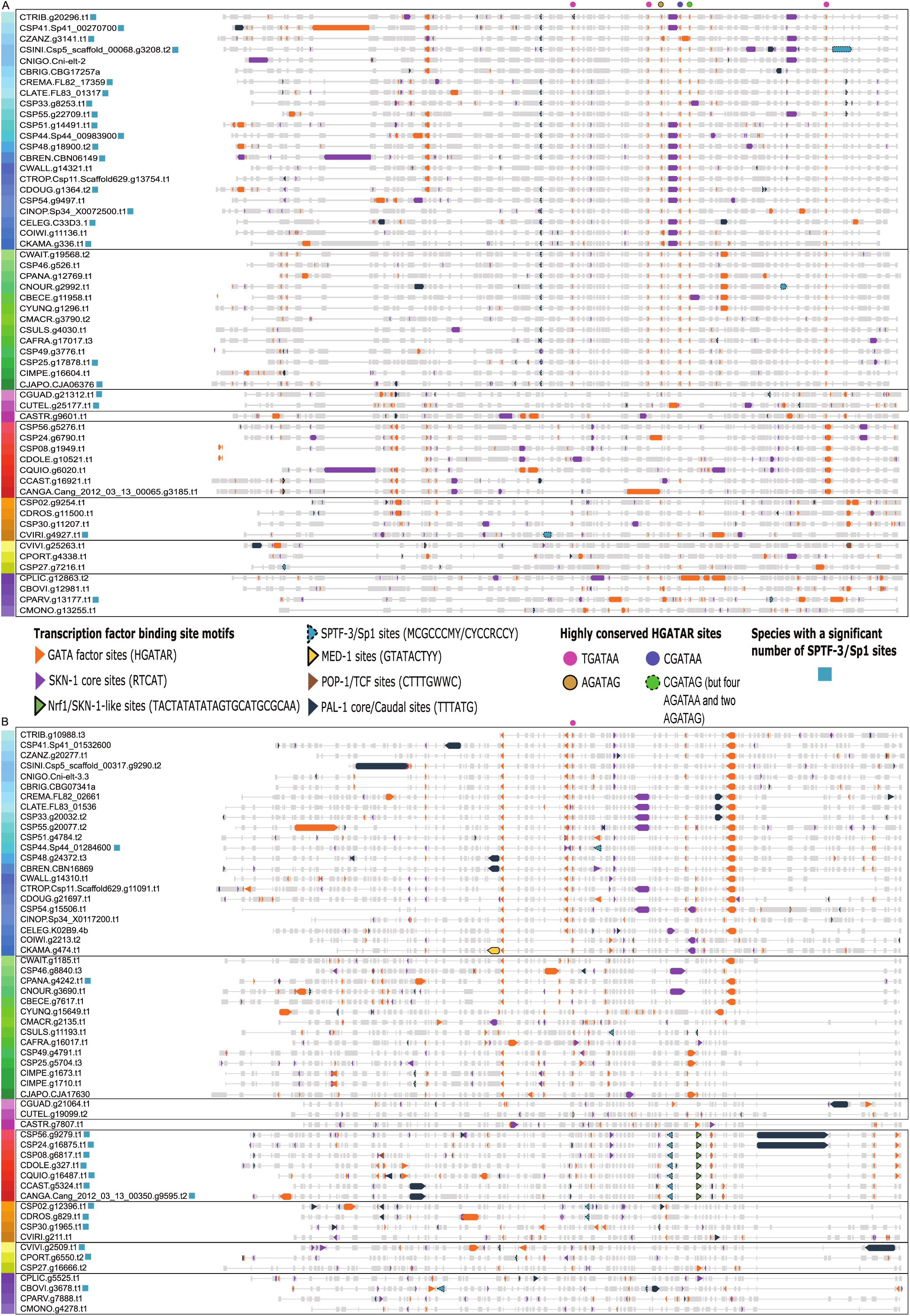
Comparison of transcription factor binding sites in *Caenorhabditis elt-3* and *elt-2* promoters. Transcription factor binding sites of interest, including those found significantly more than expected by chance, are indicated in the predicted proximal promoters of the *elt-3* (A) and *elt-2* (B) orthologs from the *Caenorhabditis* species included in this study. Aligned promoter sequences are represented by gray boxes, whereas gray horizontal lines between the boxes represent gaps in the alignment. Each entry represents the predicted proximal promoter sequence of an *elt-3* (A) or *elt-2* (B) ortholog and they are listed in the same order (top to bottom) as the *Caenorhabditis* species in the phylogeny shown in Figure 2B (left to right). The black boxes delineate the different species clades. The keys to the different transcription factor binding site motifs (depicted using triangles of different colors), and the highly conserved HGATAR sites (depicted using circles of different colors), are shown between panels (A) and (B). **(A)** *elt-3* ortholog promoter sequences. Note the highly conserved HGATAR site in the *Elegans* group species (indicated above the panel). **(B)** *elt-2* ortholog promoter sequences. Note the highly conserved HGATAR sites (colored circles) in the *Elegans* supergroup species (as highlighted above each panel).

### Elegans supergroup elt-2 orthologs may be regulated by an Sp1 family transcription factor, SPTF-3

We found significant numbers of Sp1-like (CYCCRCCY (Saito et al. 2013)) and/or SPTF-3 (MCGCCCMY (Narasimhan et al. 2015)) binding sites in the promoters of many (18 of 35) *Elegans* supergroup *elt-2* orthologs (blue squares next to gene names in Fig. 5A). SPTF-3 is a homolog of the Sp1 family in *C. elegans* (Ulm et al. 2011). Moreover, we found that a Sp1/SPTF-3 motif aligns near the middle of the putative promoters of 30 of 35 *Elegans* supergroup *elt-2* orthologs (Fig. 5A). Additionally, using MEME (Bailey and Elkan 1994), an Sp1-like motif was identified in 34 of 35 *Elegans* supergroup *elt-2* promoters, a significant hit rate (data not shown). This suggests that SPTF-3 or another Sp1 transcription factor may directly regulate *Elegans* supergroup *elt-2* orthologs.

### non-Elegans supergroup and non-Guadeloupensis group elt-3 orthologs may be regulated by a Sp1 family transcription factor, SPTF-3

To look for clues as to how *elt-3* was regulated before its expansion (see above), we searched for conserved transcription factor binding sites in the promoters of *elt-3* orthologs (see Materials and Methods). We found significant numbers of Sp1-like (CYCCRCCY (Saito et al. 2013)) and/or SPTF-3 (MCGCCCMY (Narasimhan et al. 2015)) binding sites in the promoters of most (13 of 19) *elt-3* orthologs of non-*Elegans* supergroup and non-*Guadeloupensis* group species (blue squares next to gene names in Fig. 5B). Moreover, an Sp1-like motif was the top hit identified using MEME (Bailey and Elkan 1994), which identified a similar motif in 17 of 19 *elt-3* promoters in non-*Elegans* supergroup and non-*Guadeloupensis* group species. On the other hand, similar motifs were identified in the *elt-3* promoters of only five of 37 *Elegans* supergroup and *Guadeloupensis* group species (Fig. 5B). RNAi knockdown of *sptf-3* in *C. elegans* leads to reduced *end-3* and *end-1* reporter expression and incorrectly specified endoderm (Sullivan-Brown et al. 2016). Sp1-like binding sites are also found in the promoters of most *med, end-1*, and *end-3* orthologs (Maduro 2020). Moreover, whole-embryo single-cell RNA sequencing indicates that *C. angaria sptf-3* is expressed during early embryogenesis (; Macchietto et al. 2017).

### Angaria group elt-3 orthologs may be regulated by SKN-1 orthologs

In addition to Sp1-like binding sites, SKN-1 binding sites are overrepresented in most *med*, *end-1*, and *end-3* promoters (Zhu et al. 1997; Maduro 2020). The SKN-1 orthologs in *C. elegans* and *C. briggsae* contribute extensively to initiating the endoderm specification network, primarily by activating *med-1* and *med-2* and possibly by directly activating *end-3* (Bowerman et al. 1992; Maduro et al. 2001; Maduro, Kasmir, et al. 2005a; Maduro et al. 2007; Lin et al. 2009). We found at least one SKN-1 core binding site (RTCAT; Blackwell et al. 1994) in every *elt-3* promoter we examined (Fig. 5B); however, none of them have more SKN-1 sites than expected by chance. We did not identify any additional strongly conserved transcription factor binding sites in the promoters of any of the *elt-3* orthologs, such as POP-1 or PAL-1 sites which have both been found to contribute to endoderm specification initiation in *C. elegans* (Maduro, Kasmir, et al. 2005b; Maduro et al. 2015). However, we did identify an invariant motif, TACTATATATAGTGCATGCGCAA, in the promoters of all seven *elt-3* orthologs in the *Angaria* group (Fig. 5B). We then searched the JAPSPAR 2018 core non-redundant database (jaspar.-genereg.net) for similar motifs. *Arabidopsis thaliana* FUS3, a B3 DNA-binding domain (DBD) protein, was the top hit, presumably because it binds to GCATGC; however, B3 DBDs are known to be plant-specific (Yang et al. 2021). The next best match to this invariant *Angaria* group *elt-3* motif was the *Homo sapiens* Nrf1 site: GCGCNTGCGC (jaspar.genereg.net). A BLASTp search (e-value cutoff of 0.01) did not reveal any highly conserved Nrf1 orthologs in any of the *Caenorhabditis* species included in this study. However, Nrf1 contains a bZIP DBD, and the *C. elegans* transcription factor SKN-1 also contains the basic region of a bZIP domain. Moreover, the invariant *Angaria* motif starts with a TATA-rich region, and the *C. elegans* SKN-1 DBD also contains part of a homeo-like domain which binds T/A-rich sequences (Blackwell et al. 1994; Carroll et al. 1997; Pal et al. 1997; Lo et al. 1998). Even though the *C. elegans* SKN-1 bZIP-like domain binds RTCAT sequences with high affinity (1 nM; Blackwell et al. 1994), and this exact sequence is not found in the invariant *Angaria* group motif, the specificity of SKN-1 in *C. elegans* may have diverged from that of SKN-1 in other *Caenorhabditis* species; alternatively, this invariant motif could be a secondary binding site for SKN-1 orthologs.

### Structures, motifs, and locations of the elt-3 paralogs hint at their evolutionary history

Although our data indicate that *end-1, end-3,* and *elt-7* all evolved from an ancestral *elt-3,* it is not clear whether this ancestral *elt-3* duplicated once to produce an *elt-7/end* ancestor or whether two separate duplications produced the *elt-7* and the *end-3/1* subclades (Fig. 6A vs. B). To investigate this, we examined the locations and structures of, and amino acids encoded by, all of these genes. The median protein length of ELT-3 orthologs used in this study is 322 residues, substantially longer than the median lengths of ELT-7, END-1, and END-3 proteins, which are more similar to each other (202, 226, and 240 residues, respectively). Ancestral structures of the *Elegans* supergroup *end-1*, *end-3*, *elt-7*, and *elt-3* genes predicted from the structures of extant orthologs (Darragh AC, Rifkin SA, unpublished data, https://doi.org/10.1101/2022.05.20.492891, last accessed May 23, 2022) indicate that most of these genes in this clade have an intron located at the same position in their zinc finger (ZnF) coding sequence, an intron location also found in most *elt-2* orthologs and in *Japonica* group *med* orthologs (Eurmsirilerd and Maduro 2020; Maduro 2020). Moreover, the last two exons of *Elegans* supergroup *end-1*, *end-3*, *elt-7*, and *elt-3* orthologs code for their GATA domains. The combination of this conserved intron position and the GATA domain location is only found in this clade and among the *Japonica* group *meds* (Darragh AC, Rifkin SA, unpublished data, https://doi.org/10.1101/2022.05.20.492891, last accessed May 23, 2022). Most *end-1* and *elt-7* orthologs have four exons, while most *end-3* and *elt-3* orthologs have three and eight, respectively. Nevertheless, we predict that the *Elegans* supergroup ancestral *end-3* gene was comprised of four exons because of the conservation of a poly-serine motif at the N-terminus of the END proteins, the fact that the ZnFs are coded for in the last two exons of *end-1* and *end-3*, and the many *end-3* orthologs that have four exons (Maduro 2020); all this evidence is consistent with a full-length duplication of an ancestral *end* gene producing the ancestral *end-1* and *end-3* genes.

**Figure 6.**
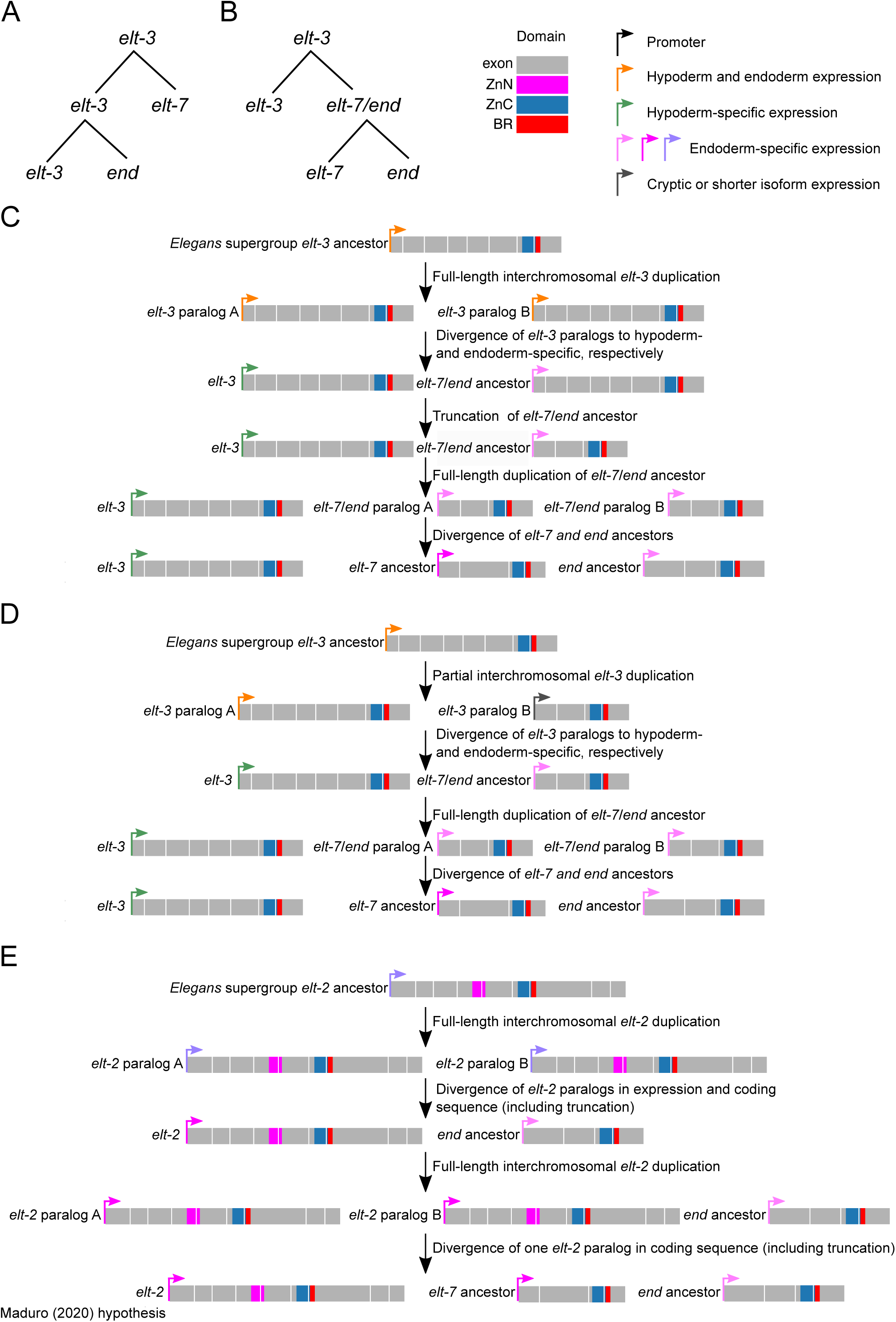
Scenarios for how initiation of the expansion of endoderm specification GATA factors could have occurred. Comparison of possible gene duplication scenarios for initiating GATA factor expansion, those supported by our results (A-D) and another proposed by Maduro (Maduro 2020) (E). **(A)** Scenario involving two duplications of *elt-3,* one which produced the ancestor of *elt-7* and another which produced the ancestor *end* gene. **(B)** Scenario involving a single *elt-3* duplication, in which one duplication of *elt-3* produced the ancestor *elt-7/end* gene and then a subsequent duplication of the *elt-7/end* ancestral gene produced the ancestors of the *elt-7* and *end* genes. **(C)** Details of the proposed scenario involving a single duplication of a full-length *elt-3*. (Alternatively, if instead of one, two full-length *elt-3* duplications occurred, then the first three steps of this scenario could occur twice to produce the *elt-7* and *end* ancestor genes.) **(D)** Details of a proposed scenario involving a single, partial duplication of *elt-3*. (Alternatively, if instead of one, two partial-length *elt-3* duplications occurred, the first two steps of this scenario could occur twice to produce the *elt-7* and *end* ancestor genes.) **(E)** Molecular representation of a previously published hypothesis (Maduro 2020) for how two *elt-2* duplications could have produced the *elt-7* and *end* ancestor genes. The key to color-coding of gene domains and expression patterns is located in the upper right corner of the figure.

We found no instances in which an *elt-3* ortholog occurred on the same piece of genomic DNA as an *elt-7*, *end-1*, or *end-3* ortholog. In fact, in the six species for which chromosome-level assemblies were available, *elt-3* orthologs are found on the X chromosome, while *elt-7*, *end-1*, or *end-3* orthologs are found on chromosome 5. Based on an analysis of synteny (see Materials and Methods), these locations are likely consistent throughout the genus. Moreover, *elt-7*, *end-1*, and *end-3* orthologs are syntenic in one additional species. All of these pieces of evidence suggest that the *elt-7* and the *end* genes share a more recent history with each other than with *elt-3* (i.e., Fig. 6B). An additional piece of evidence is more equivocal. Maduro and colleagues(Maduro, Hill, et al. 2005; Maduro 2020) identified a poly-serine motif near the N-terminus of the END-1 and END-3 orthologs they examined. We found such a motif in 30 of 35 ELT-3 orthologs in the *Elegans* supergroup and four of 23 ELT-3 orthologs in the non-*Elegans* supergroup; on the other hand, we only found this motif in three of the 35 *elt-7* orthologs we examined. Other than their GATA domains and this poly-serine motif, we found no additional sequence homology among the ELT-3, ELT-7, and the END orthologs.

### Evidence of relaxed selection on one paralog relative to the other

Because gene duplication changes the functional context of genes we tested whether the intensity of selection changed after the *elt-3* expansion using RELAX (Wertheim et al. 2015). Our results indicated that in the *Elegans* supergroup both the *elt-7* orthologs (p<0.0001; k=0.34) and the *end* orthologs (p<0.0001; k=0.44) experienced less intense selection than did the *elt-3*s. In turn, selection intensity relaxed on the *end-3* ortholog group after duplication as compared to the *end-1* orthologs (p<0.0001; k=0.47). Additionally, selection on *Elegans* supergroup *elt-3*s has intensified since expansion as compared to selection on non-*Elegans* supergroup *elt-3*s (p<0.0001; k=2.64). All of these patterns of selection intensity are concordant with the differences in branch lengths among these groups that are readily apparent in our phylogenetic tree (Fig. 2A).

### The evolutionary history of med orthologs is opaque due to quick turnover

While our phylogenetic reconstruction supports a clear hypothesis about the origin of *end-3, end-1,* and *elt-7*, the *med* orthologs sit in a subclade of their own, with no clear phylogenetic connection to other groups (Fig. 2A). *med* genes code for the shortest GATA-domain-containing proteins that we identified in the 60 species included in our study (Darragh AC, Rifkin SA, unpublished data, https://doi.org/10.1101/2022.05.20.492891, last accessed May 23, 2022). The *Elegans* group *meds* have no introns, while those in the *Japonica* group have one to three introns, including one splice site in the same location in their ZnF as is found in *elt-3* and its paralogs and in the *elt-2* orthologs (Maduro 2020), but which is not found in any of the other GATA-domain-containing proteins we identified (Dar - ragh AC, Rifkin SA, unpublished data, https://doi.org/10.1101/2022.05.20.492891, last accessed May 23, 2022). Although the *C. elegans* MEDs bind an atypical motif (Broitman-Maduro et al. 2005; Lowry et al. 2009), their DNA-binding domains more closely resemble canonical GATA domains than they do the atypical GATA domains of EGL-27, SPR-1, or RCOR-1 (Darragh AC, Rifkin SA, unpublished data, https://doi.org/10.1101/2022.05.20.492891, last accessed May 23, 2022), supporting the hypothesis that the MEDs arose from one of the canonical *Caenorhabditis* GATA factors instead of from a different, more atypical GATA-domain-containing protein. Syntenic *med* paralogs are usually relatively close to each other (range of 319 bp to 13 kb, median of 3.1 kb) and similar in sequence (93% median nucleo - tide identity, compared to 77% for non-syntenic *med* paralog pairs). However, most sister species have *med* paralogs on at least two different chromosomes. Our phylogeny (Fig. 2A) shows that most MEDs are most closely related to paralogs within their own species and have highly variable copy numbers across species, indicating that these genes evolve through rapid duplication and loss (Maduro 2020).

## Discussion

Our analysis of the evolution of GATA factors in the 58 *Caenorhabditis* species for which protein sequences are currently available shows that the genes of most of the GATA factors involved in endoderm development – *end-1*, *end-3*, *elt-7*, and the *med* genes – arose during the course of a larger GATA factor expansion in the genus (Fig. 2). Although this radiation re-wired the endoderm specification network, it was not associated with any known change in the environment, morphogenesis, or anatomy of these animals. Additionally, we found that the role of the most downstream GATA factor in this network, encoded by *elt-2*, is likely conserved across the genus (Fig. 3A,C; Fig. 4). Interestingly, five GATA factor binding sites were likely fixed in *elt-2* promoters at the base of the *Elegans* supergroup concurrent with the elaboration of its *trans*-activating network (highlighted by the colored dots at the top of Figure 5A). The concentration of a single type of transcription factor in a gene regulatory network – especially one as temporally and spatially restricted as the endoderm network – is extraordinarily rare and creates the potential for extensive developmental system drift.

Maduro (Maduro 2020) hypothesized that two *elt-2* duplications in the *Elegans* supergroup ancestor produced an ancestral *end/med* gene and an ancestral *elt-7* gene (Fig. 6E). This hypothesis was supported by the fact that *elt-2*, *elt-7*, *end-1*, *end-3*, and the *med* orthologs all function in the *C. elegans* endoderm specification network (Zhu et al. 1997; Broitman-Maduro et al. 2005; McGhee et al. 2007; McGhee et al. 2009; Sommermann et al. 2010; Dineen et al. 2018), *elt-2* orthologs are conserved throughout the *Caenorhabditis* genus, and *elt-2* orthologs share a conserved intron location within their ZnFs with *end-1*, *end-3*, *elt-7*, and *Japonica* group *meds*. However, *elt-3* orthologs have the same conserved intron location in their ZnFs which has been conserved in all 58 *Caenorhabditis* species we examined (Darragh AC, Rifkin SA, unpublished data, https://doi.org/10.1101/2022.05.20.492891, last accessed May 23, 2022). Moreover, our phylogeny indicates that ELT-3 orthologs share a more recent common ancestor with ELT-7, END-1, and END-3 orthologs than with any other *Caenorhabditis* GATA-domain-containing orthologs (Fig. 2A), and our smFISH analysis indicates that *C. angaria elt-3* mRNA is expressed in the early endoderm (Fig. 4B,C). Based on this new evidence, we argue that one or two *elt-3* duplications in the *Elegans* supergroup ancestor produced the ancestors of the *end* and *elt-7* genes.

Consistent with previous results (Maduro, Hill, et al. 2005; Boeck et al. 2011; Maduro 2020), our phylogeny (Fig. 2A) indicates that END-3 and END-1 orthologs share a recent common ancestor; the evolutionary relationship(s) between ELT-3, ELT-7, and the END ancestor is less clear, however. The ELT-7 ortholog group branches off of a shared node between the ELT-3 and the END orthologs (0.2 substitutions per site in Figure 2A), suggesting that END orthologs are more closely related to ELT-3 orthologs than to ELT-7 orthologs. This finding supports a scenario in which separate *elt-3* duplications produced the *elt-7* and *end* ancestors, as opposed to a single *elt-3* duplication producing the ancestor of both the *elt-7* and *end* genes followed by a second duplication of this *elt-7*/*end* gene (Fig. 6A vs. B). However, both the structures and chromosome locations of these GATA-factor-encoding genes support a scenario in which a single *elt-3* duplication (along with a single shortening and interchromosomal rearrangement event) gave rise to the *elt-7/end* ancestor (e.g., Fig. 6C or D). If the ELT-3 and END N-terminal poly-serine motifs and/or the SPTF-3 regulatory sites in the promoters of the non-*Elegans* supergroup and non-*Guadeloupensis* group *elt-3* orthologs and *end* genes are homologous, it supports a scenario in which a full-length duplication of the *elt-3* coding sequence produced the *end* ancestor (followed by sequence loss to produce the final four-exon version). If the *elt-7* and *end* ancestors arose from the same *elt-3* duplication, then their ancestor likely quickly duplicated again since these ortholog groups experienced different subsequent trajectories of mutation and deletion. However, if the *elt-7* and *end* ancestors resulted from two different *elt-3* duplications, then the lack of a poly-serine motif and SPTF-3 binding sites in *elt-7* orthologs we observed (data not shown) could be the result of a partial *elt-3* duplication producing the ancestral *elt-7* gene or due to greater relaxation of selection pressure experienced by *elt-7* orthologs. This ambiguity in the precise birth order of the *elt-7* and *end* gene ancestors may reflect the fact that this radiation happened in an evolutionarily short period of time such that both *elt-7* and *end* orthologs are about equally diverged from *elt-3* orthologs, albeit in different ways.

The expression of *C. elegans end-3* starts before that of *end-1*, whereas the peaks of *end-3* and *end-1* mRNA expression occur in first (1E) and second (2E) cell stages of endoderm development, respectively (Raj et al. 2010). *C. elegans elt-7* expression starts after that of *end-1*, during 2E (Sommermann et al. 2010). *C. elegans elt-7* continues to be expressed for the lifetime of the worm, whereas the *ends* genes are only expressed transiently during endoderm specification (Raj et al. 2010; Sommermann et al. 2010). Based on our finding that *C. angaria elt-3* is expressed similarly to *C. elegans end-1* in the endoderm (Fig. 3B-D), we predict that this was the endoderm expression pattern of the *Elegans* supergroup ancestral *elt-3* paralog. This suggests that the expression patterns of *end-3* and *elt-7* diverged from that of their ancestor. Despite this apparent divergence in gene expression patterns, the functions of *C. elegans end-3*, *end-1*, and *elt-7* have been found to be interchangeable. For example, the expression of *end-3*, or *end-1*, or *elt-7* in the early endoderm is sufficient for gut specification (Zhu et al. 1998; Maduro, Hill, et al. 2005; Wiesenfahrt et al. 2016) and ectopic expression of any of these GATA factors is sufficient to activate expression of endoderm markers (Maduro and Rothman 2002; Sommermann et al. 2010). This suggests that these paralogs have primarily functionally diverged through *cis*-regulatory changes, while their protein sequence differences have not been found to have functional consequences. However, the DNA binding preferences of *C. elegans* END-3 and END-1 are less specific than those of *C. elegans* ELT-7 (Narasimhan et al. 2015). The ENDs bind to GATA sequences, without much preference regarding the flanking base pairs, while ELT-7 prefers to bind to TGATAA sequences (Narasimhan et al. 2015). Moreover, we found that non-TGATAA HGATAR sites are highly conserved in *Elegans* supergroup *elt-2* promoters (Fig. 5A), suggesting that these non-TGATAA sites are more likely bound by the ENDs while the TGATAA sites are preferably bound by ELT-7 and ELT-2. Therefore, the binding preference of ELT-3 paralogs may have diverged in parallel with the expression pattern of *elt-3*.

Given the resemblance of the invariant Nrf1/bZIP motif in *Angaria* group *elt-3* promoters to a possible SKN-1 binding site (see Results) and the involvement of SKN-1 in the endoderm specification network – at least in *Elegans* supergroup species (Maduro 2020), we hypothesize that *Angaria* group SKN-1 orthologs bind to this invariant sequence to activate *elt-3* expression in early endoderm cells (Fig. 7 left side). If true, and if ELT-3 is indeed part of the endoderm specification network in non-*Elegans* supergroup species, then any regulation of the network involving SKN-1 should be conserved in the initial stages of endoderm specification, despite change in the SKN-1 binding site.

**Figure 7.**
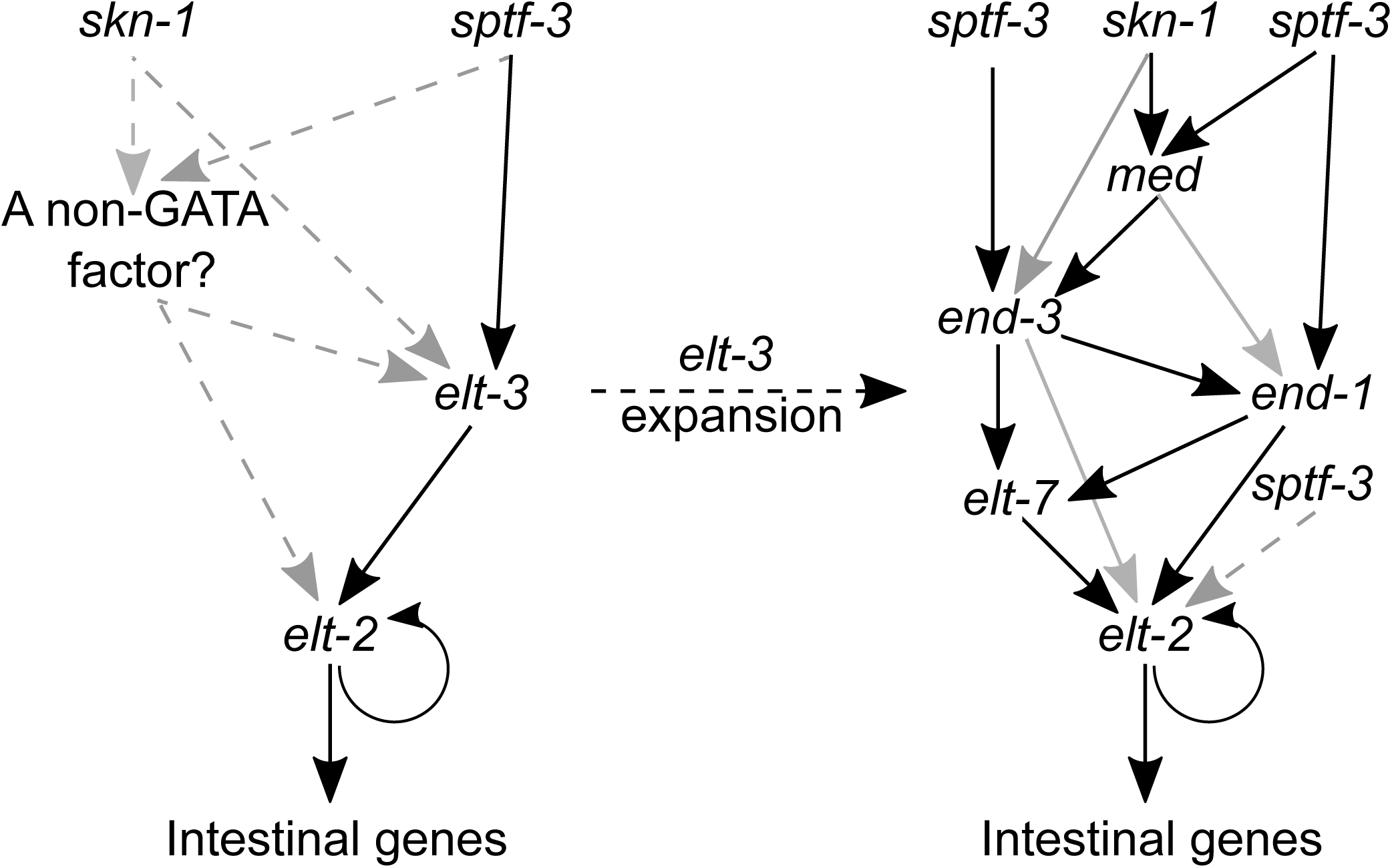
Evolutionary model of how GATA factors expanded in the endoderm specification network. Data from this study are consistent with this evolutionary model in which, prior to our proposed expansion of the *elt-3* gene in the endoderm specification network (left side of figure), the functioning of this network was initiated by expression of *sptf-3* and/or *skn-1*, which activated *elt-3* (and possibly another transcription factor expressed earlier, depicted as “A non-GATA factor?”). Expression of ELT-3 (and possibly other transcription factors) then activated *elt-2*. ELT-2 then likely regulated hundreds of genes expressed specifically (or primarily) in the intestine and perhaps auto-regulated its own gene expression. This “pre-expansion” network (shown on the left) is expected to be similar to the endoderm specification networks found in non-*Elegans* supergroup and non-*Guadeloupensis* group species, like *C. angaria*. Our data suggest that a duplication(s) of *elt-3* led to the addition of three or four GATA factor paralogs to the endoderm specification network that function between *sptf-3* and/or *skn-1* and *elt-2* resulting in the network shown on the right. This model predicts that during the GATA factor expansion *elt-3* paralogs subfunctionalized into: an *elt-3*-like gene expressed only in the hypoderm (not shown), an endoderm-specifically expressed *elt-7*, and an ancestor of the *end* genes. (See Figure 6A-D for molecular details of how this subfunctionalization could have occurred). Data from this study also support the previously proposed hypotheses that an additional *end* gene duplication produced the ancestors of *end-1* and *end-3* (Maduro, Hill, et al. 2005; Coroian et al. 2006) and that another *end* gene duplication likely produced the ancestor *med* gene (Maduro 2020). Neither we nor Maduro (Maduro 2020) found POP-1 nor PAL-1 transcription factor binding sites overrepresented in *end-1* (or *end-3*) promoters and therefore they are not included in the network on the right. Black arrows indicate well supported regulatory connections, while gray and dashed gray arrows represent weaker and not as well supported interactions, respectively.

While our phylogeny strongly suggests that an *elt-3* expansion occurred in the *Elegans* supergroup ancestor (Fig. 2A), the tree also suggests that additional *elt-3* duplications occurred elsewhere in the genus. We identified divergent *elt-3* paralogs in at least two of the three *Guadeloupensis* group species and in *C. astrocarya*, a species likely basal to the *Elegans*/*Guadeloupensis* species (Fig. 2); interestingly, most of these divergent *elt-3* paralogs have shorter gene structures (two to six exons and median of 220 amino acids) more like those of *elt-7*, *end-1*, and *end-3* (Darragh AC, Rifkin SA, unpublished data, https://doi.org/10.1101/2022.05.20.492891, last accessed May 23, 2022) and, like the *Elegans* supergroup *elt-3*s, their promoters do not have SPTF-3 binding motifs (Fig. 5B). Therefore, our evidence also fits the alternative scenario that *elt-3* duplication, shortening, and subfunctionalization occurred one stage earlier, i.e. in the *Elegans*/*Guadeloupensis*/*astrocarya* ancestor, followed by extensive divergence of *elt-3* paralogs in the different lineages.

In *C. elegans, elt-2* overexpressed under the control of the *end-1* promoter can compensate for loss of *end-3*, *end-1*, *elt-7*, and *elt-4* (Wiesenfahrt et al. 2016). Analogous expression of *C. elegans elt-3* cannot do this. This is especially surprising considering that ELT-3 orthologs are more closely related to END-3, END-1, and ELT-7 orthologs than to ELT-2 orthologs (Fig. 2A). This suggests that *C. elegans* ELT-3 (and likely all *Elegans* supergroup ELT-3 orthologs) has lost the ability to specify endoderm even when ectopically expressed there and even though it can bind TGATAA sites (Narasimhan et al. 2015). However, our finding that mRNA of the *C. angaria elt-3* ortholog is expressed in early endoderm cells (Fig. 3B,C) in a pattern reminiscent of *end-1* expression, suggests that it probably also functions there. Not only did the *Elegans* supergroup ancestral *elt-3* likely subfunctionalize its expression pattern, something else must have changed about the coding region in its descendants such that one branch preserved its capacity to function in the endoderm while the other branch lost this ability. We did not find any obvious differences between the DNA binding domains in the *elt-3* orthologs in *Elegans* supergroup versus non-*Elegans* supergroup species. However, we did find a highly conserved, possible SUMOylation site (Chang et al. 2018) towards the N-terminus of most (19 of 23) ELT-3 orthologs in non-*Elegans* supergroup species, and that this [VIA]KE[ED] motif has been lost from all the ELT-3 orthologs in *Elegans* supergroup species (Darragh AC, Rifkin SA, unpublished data, https://doi.org/10.1101/2022.05.20.492891, last accessed May 23, 2022). We therefore speculate that ELT-3 orthologs in non-*Elegans* supergroup species could undergo post-translational modification (associated with this putative SUMOylation site) and be involved in an endoderm-specific protein-protein interaction(s). Interestingly, our finding that selection has been more intense on the *elt-3* orthologs in *Elegans* supergroup species compared to those in the non-*Elegans* supergroup species, may be reflective of a functional change in the *Elegans* supergroup ELT-3s.

We found no additional evidence for a *med* gene ancestor originating from an *elt-2* duplication (as proposed by Maduro (Maduro 2020)) but, rather, more evidence that the *med* ancestor originated from a duplication within the *end-3/end-1/elt-7/elt-3* clade in the *Elegans* supergroup ancestor. The many species-specific paralogs and long phylogenetic branches found in the MED ortholog group (Fig. 2) suggest that the *med* genes turn over quickly, as previously noted (Maduro 2020). This quick turnover has likely erased additional evidence relating to the relationship between the MED orthologs and other GATA factors. The strongest evidence for the origin of the *med* orthologs (on which Maduro (Maduro 2020) based his hypothesis) is the location of an intron in the ZnF of most *Japonica* group *meds* (that has been lost from *Elegans* group *meds*), which is found at the same location in the ZnFs of *end-1*, *end-3*, *elt-2*, and *elt-7* orthologs as well as of *elt-3* orthologs, as we have shown (Darragh AC, Rifkin SA, unpublished data, https://doi.org/10.1101/2022.05.20.492891, last accessed May 23, 2022). The structures of *Japonica* group *med* genes are also most similar to those of *end-3* homologs (Darragh AC, Rifkin SA, unpublished data, https://doi.org/10.1101/2022.05.20.492891, last accessed May 23, 2022). Since the *Elegans* supergroup ancestral *end-3* gene may have lost an intron after diverging from *end-1* (see Results), a full-length *end-3* duplication could have produced the *med* ancestor. However, a partial duplication of an *end-1*, *elt-2*, *elt-3*, or *elt-7* ortholog, as well as a full duplication of any of these ancestral genes followed by coding sequence loss, are additional possible explanations. Since we have shown that three gene duplications occurred within the *end-3/end-1/elt-7/elt-3* clade and were likely fixed in the *Elegans* supergroup ancestor (Fig. 2), it is plausible that at least one more could have occurred to produce the *med* clade.

In conclusion, our data suggest that *elt-2* plays a consistent role throughout the *Caenorhabditis* genus in regulating, through TGATAA binding sites, hundreds of genes expressed specifically or predominantly in intestine (Fig. 3A,C,D; Fig. 4; Fig. 7). As we have shown in *C. angaria* (Fig. 3B-D; Fig. 5; Fig. 7), we predict that *elt-3* orthologs function before *elt-2* orthologs in the endoderm specification network of non-*Elegans* supergroup species and did so in the *Elegans* supergroup ancestor. Evidence also suggests that the *Elegans* supergroup ancestral network may have been initiated by SPTF-3 activating *elt-3* (Fig. 5B). SKN-1 may also play a conserved role in the initiation of this network across the genus, through an as yet unknown gene(s) upstream of *elt-3* that is analogous to the *med* and *end-3* genes which were subsequently displaced by the radiating GATA factors (Fig. 7). Or SKN-1 may directly regulate *elt-3*, even though we did not find significant numbers of SKN-1 binding sites in *elt-3* promoters (Fig. 5B; Fig. 7). Figure 7 summarizes our proposed model of how the *Elegans* supergroup ancestral endoderm specification network evolved in a relatively short period of time, all without any apparent phenotypic change.

## Materials and Methods

### Phylogenetic analysis

GATA factor homolog identification and an initial phylogenetic analysis was done for a companion paper (Darragh AC, Rifkin SA, unpublished data, https://doi.org/10.1101/2022.05.20.492891, last accessed May 23, 2022). We used the same alignment and tree building procedure, with an additional 3000 ultrafast bootstraps (Minh et al. 2013; Hoang et al. 2018), for creating a phylogenetic inference of the 714 protein sequences we deemed “confident” (see Darragh AC, Rifkin SA, unpublished data, https://doi.org/10.1101/2022.05.20.492891, last accessed May 23, 2022) for the protocol used to determine “confident” proteins); the resulting maximum likelihood tree is shown in Figure 2A.

### Process for classifying proteins as singletons, paralogs, representative paralogs, or divergent paralogs

We classified the 714 proteins we were most confident in as a *singleton* or *paralog*, depending on how many proteins from each species clustered into the 12 ortholog groups in our phylogeny (Fig. 2A). For example, a species with only a single ELT-3-like sequence grouping in the ELT-3 ortholog group was considered a singleton ELT-3 ortholog whereas a species with multiple ELT-3-like sequences that robustly grouped into the ELT-3 ortholog group were deemed paralog ELT-3s. Other than in the MED ortholog group, which was comprised primarily of paralogs, the END-3 ortholog group contained the most confident paralogs (Fig. 2A) and, since we focus here on how an *elt-3* radiation produced *end-3*, *end-1*, and *elt-7*, we further categorized these paralogs as either *representative* or *divergent*. This classification is based on how conserved the paralog’s sequence is in relation to the sequences of singletons in the same ortholog group; if an individual paralog within a species had a noticeably higher level of conservation to singleton orthologs than did other paralogs, we choose that paralog as the representative one and labeled the others divergent. If multiple paralogs within a species exhibited approximately equal levels of conservation to singleton orthologs, they were all considered representative paralogs. For most analyses, we used both representative paralogs and singletons; for most analyses of the MED ortholog group, we used single ZnF paralogs and singletons; and for the other groups we used only singletons for most analyses.

### Worm maintenance

*C. angaria* strain PS1010 was grown at room temperature (RT, approximately 21-22 °C) on Nematode Growth Medium Lite (NGM Lite, 34.22 mM NaCl, 4g/L Bactotryptone, 22.04 mM KH_2_PO_4_, 2.87 mM K_2_HPO_4_, 20.69 uM Cholesterol, 59.47 mM Agar, in distilled (DI) water) in Petri dishes containing a lawn of *Escherichia coli* strain OP50 as a food source, in a manner similar to that standardly used for culturing and maintaining *C. elegans* (Brenner 1974; Stiernagle 2006).

### Single molecule fluorescence in situ hybridization

*C. angaria* strain PS1010 embryos were isolated from gravid adults using worm-bleaching solution (250 mM NaOH, 1% NaOCl, in DI water) and then, following standard *C. elegans* protocols for synchronizing them, grown in liquid M9 (22 mM KH_2_PO_4_, 42 mM Na_2_HPO_4_, 85.6 mM NaCl, 1 mM MgSO4, in DI water) for one day until they hatched. Synchronized, larval stage one (L1) worms were then pipetted onto NGM Lite agar plates with *E. coli* lawns and grown, using standard procedures, at RT for four days until the by then adult worms started laying eggs. To enrich for early embryos (i.e., those still inside the worms), the *C. angaria* on the plates were washed off with DI water and into a 40 um filter setup which retained adults and let already laid eggs pass through to be discarded. The adult worms were then treated with worm-bleaching solution (as described above) to extract early embryos (Raj et al. 2008). Embryos were then fixed with 4% formaldehyde in PBS (137 mM NaCl, 2.7 mM KCl, 8 mM Na_2_HPO_4_, 2 mM KH_2_PO_4_ in nuclease-free water), freeze-thawed using liquid nitrogen, washed with 1x PBS, placed in 70% ethanol in nuclease-free water (Ambion), and stored at 4 °C for at least overnight and up to one week. Embryos were then washed in a solution (wash buffer) comprised of 10% formamide, 2x SSC (300 mM NaCl, 30 mM Na_3_C_6_H_5_O_7_) prepared using nuclease-free water. Hybridizations were carried out as previously described (Raj et al. 2008; Raj et al. 2010) in 100 uL hybridization buffer (10% formamide, 2x SSC, 0.1g/mL dextran sulfate in nuclease-free water) to which 1 ul of each of two smFISH probes, one designed to hybridize to *elt-2* mRNA (Atto 647::*elt-2*, Biosearch) and the other to *elt-3* mRNA (Quasar 570::*elt-3*, Biosearch), had been added; embryos were incubated in the hybridization solution for 16 hours at 30 °C. Embryos were then washed with wash buffer and their nuclei stained with 5 ug/mL DAPI (4′,6-diamidino-2-phenylindole, Roche) prepared in wash buffer for 10 minutes at 30 °C.

For imaging, embryos were suspended in RT glox buffer comprised of 20 mM Tris Cl pH 7.5, 2x SSC, 0.4% glucose in nuclease-free water, 37 ug/mL glucose oxidase, and 1 ul catalase (Raj et al. 2010; Sigma-Aldrich). Embryos were imaged in Z-stacks with 0.3 um spacing at 100x magnification on a Nikon epifluorescent compound microscope. smFISH signals were quantified using a machine-learning spot-classification tool, AroSpotFinding Suite (Rifkin 2011; Wu and Rifkin 2015), and visually con-firmed. Nuclei were counted manually with the help of MATLAB. An embryo’s nuclei count was used as a proxy for its developmental stage.

### Identifying TGATAA sites in the promoters of orthologous genes expressed gut-, muscle-, neural-, or hypoderm-specifically/enriched

We identified the longest isoform among each of the following groups of *C. elegans* genes that are specifically expressed or enriched for expression (as per McGhee et al. 2007, 2009) in the organ/ tissue indicated: 197 intestine, 71 muscle, 47 neural, and 79 hypoderm. We then used reciprocal BLASTp (e-value cutoff 0.01) to search for putative orthologs of these genes in the 57 available non-*C. elegans Caenorhabditis* and two outgroup nematode proteomes. We only included orthologs with an ATG at the start of their coding sequence and genes that had at least two orthologs. Next, we identified the putative proximal promoters (i.e., 2 kb upstream of each coding sequence start, if available) of each ortholog. Some gene start codons are very close to the end of their scaffold/contig, if there was less than 5 bps upstream of the ATG we did not include this sequence as a promoter. We then determined the number of TGATAA sites in each promoter. Next, we used hierarchical clustering with Euclidean distance metrics to organize the genes by number of TGATAA sites in their promoters (and whether the species had a putative ortholog). These data are what is plotted in Figure 4.

### Identifying conserved transcription factor binding sites in elt-3 and elt-2 promoters

We extracted the sequences comprising 1200 bps upstream of the start codons (i.e., proximal promoters) of *elt-2* singletons and *elt-3* singletons and representative paralogs from the scaffold files of each species (see above and note that there were no confident *elt-2* paralogs). If another annotated coding sequence occurred within the upstream 1200 bps of a gene, we shortened the proximal promoter so as to eliminate the annotated coding sequences. To look for possible *cis*-regulatory motifs within these sequences we used meme-5.2.0 (Bailey and Elkan 1994) command line tools (downloaded from meme-suite.org/tools/meme) to identify any enriched motifs in the *elt-2* and *elt-3* promoters respectively. To look for clade-specific motifs we also compared *elt-2* promoters and *elt-3* promoters between the *Elegans* supergroup and non-*Elegans* supergroup species. The parameters used for our MEME analysis included consideration of both DNA strands (revcomp), motif widths between five to 12, expected site distribution, and any number of repetitions (anr, which often identified more repetitive A/T-rich motifs in *elt-3* promoters) or zero or one occurrence (zoops); the program usually stopped finding additional motifs after a significant motif with an e-value greater than 0.001 had been identified. For evaluations of *Elegans* supergroup *elt-2* promoters using the zoops option, the analysis reached the maximum number of motifs to be identified of 20.

Additionally, we looked for conserved binding sites for specific transcription factors. We aligned *elt-2* and *elt-3* promoters, respectively, using MAFFT FFT-NS-2 (Katoh et al. 2002) and searched for the following specific sites identified using MEME: canonical and non-canonical GATA-factor-DNA-binding sites, and binding sites for the *C. elegans* endoderm transcription factors SKN-1, POP-1, SPTF-3, and PAL-1 (Maduro, Kasmir, et al. 2005b; Maduro et al. 2015) (Fig. 1A). We determined whether these sites occurred in individual promoters more often than expected by chance, assuming a Poisson distribution and the sequence composition of the given promoter, in a manner similar to prior analyses carried out on previously available *end-1*, *end-3*, and *med* promoters (Maduro et al. 2015; Maduro 2020). We plotted sites in Figure 5 using Geneious 10.2.6 and Inkscape 1.0.2.

### Gene structure comparisons and predictions of ancestral gene structures

The locations of the GATA ZnF domains that we identified in each confident protein (see above) are listed in Supplemental Table 1. Eurmsirilerd and Maduro (Eurmsirilerd and Maduro 2020) defined a poly-S region as at least six serines within ten adjacent residues. Using a custom Python script, we found the locations of any such motifs and list them in Supplemental Table 1. Using the exon lengths and the domain/motif location information, we created representations of the gene structures of all the genes we deemed “confident” in this study and for which genomic data was available using a custom R script (data to be reported elsewhere). We also marked the locations of the possible SUMOylation sites ([VIA]KE[ED]; Chang et al. 2018) that we found in *elt-3* orthologs (Supp. Table 1). (Note: the *C*. sp. *45* and *C*. sp. *47* genes were excluded from this analysis because only transcriptome data were available for these species.)

We visually compared the gene structures of the confident genes (which will be reported elsewhere) and, using the principle of parsimony (and when parsimony was not sufficient to distinguish between two alternatives also treating intron loss as more frequent than intron gain (Roy and Penny 2006)), then predicted ancestral gene structures (exon number and domain location(s)) for the END-3, END-1, ELT-7, ELT-3, ELT-2, and MED ortholog groups. To estimate the lengths of the exons and introns in the ancestral genes, we calculated and used the median lengths of the exons and introns of the orthologs that had the same gene structure as the predicted ancestor.

### Predicting chromosome location

Since the genome assemblies of most of the species used in this study lack chromosome-level resolution we also used a custom Python script to identify all annotated genes within 70 kb upstream and downstream of each confident gene (“neighbor genes”). We then used BLASTp (e-value cutoff 0.1) to search for each neighbor gene’s longest isoform “tophit” in the *C. elegans* proteome and then determined which chromosome that *C. elegans* tophit was coded on.

### Testing for extent of selection pressure on paralogs and orthologs

RELAX (Wertheim et al. 2015) compares two sets of branches in a phylogenetic tree and evaluates whether the data better fits a single distribution of a few ratios of the number of non-synonymous substitutions per site to the number of synonymous substitutions per site (dN/dS) as rate categories among all branches, or different distributions for each set where the rate categories in one are related to the rate categories in the other by an exponentiation factor (k). We used RELAX with default settings to test four hypotheses about the strength of selection between sets of branches in the clades of *elt-3*, *elt-7*, *end-1*, and *end-3* orthologs. In several cases there were paralogs of the same gene within a species (e.g., two *end-3s*) for which one gene was more conserved and the other(s) more divergent. In these cases, we only included the more conserved gene in our analysis, which made our tests more conservative. We used three possible rate categories and the RELAX default settings for each test.

## Acknowledgements

We thank Lewis Stevens for pre-publication access to *Caenorhabditis* genomic data. We gratefully acknowledge The Caenorhabditis Genome Project for *Caenorhabditis* and outgroup datasets. Special thanks to Rifkin lab members for their helpful discussions. This work was supported by the National Institutes of Health (R01 GM103782); and the National Science Foundation (IOS 1936674).

